# Guiding humans towards ergonomic postures using an active exoskeleton

**DOI:** 10.1101/2025.05.14.653966

**Authors:** Waldez Gomes, Lucas Quesada, Bastien Berret, Nicolas Vignais, Dorian Verdel

**Affiliations:** Université Paris-Saclay, Inria, CIAMS, 91190 Gif-sur-Yvette, France; Enchanted Tools, 75011, Paris, France; LURPA, ENS Paris-Saclay, Université Paris-Saclay, 91190 Gif-sur-Yvette, France; Université Rennes 2, M2S, Inria, 35000 Rennes, France; Bioengineering department, Imperial College of Science, Technology and Medicine, London W12 0BZ, United-Kingdom

**Keywords:** Biofeedback, physical ergonomics, human motor control, task-relevant variability, haptic guidance

## Abstract

In the past decades, active exoskeletons have been dedicated to reducing human effort, in particular to assist workers in occupational environments. However, this approach does not promote the learning of more ergonomic postures by workers, which is critical for the long-term prevention of musculoskeletal disorders. Alternatively, we propose to use exoskeletons as biofeedback systems, generating task-relevant perturbations guiding users towards ergonomic postures. To test this approach, participants performed reach-to-hold movements towards a redundant target, allowing multiple final postures. We then introduced vibrations with posture-dependent intensity, generating a sensorimotor disturbance that canceled out either above or below each participant’s nominal preferred posture. Interestingly, participants adapted to minimize the vibrations, whether it increased or decreased the gravity efforts, and retained the novel posture when it induced lower effort. Finally, all participants significantly reduced effort post-exposure, demonstrating the feasibility and the benefits of using exoskeleton as biofeedback systems to improve posture.

## 1 Introduction

In the past decades, active exoskeletons have been used to reduce the human effort, whether it be to allow neurorehabilitation [1] or to assist workers while performing demanding tasks [2]. A potential limit of this approach is that it does not aim at guiding the user to learn new, more efficient and ergonomic postures. In fact, simply reducing the amount of human effort required to perform a task could even lead to the adoption of detrimental postures, which may shift the risk of developing musculoskeletal disorders (MSD) from a joint to another [3, 4]. The ability to simultaneously assist movements and teach efficient and protective postures to the exoskeleton’s user would be critical to reduce the long-term risk of MSD. In the present study, we investigate a method using active exoskeletons to guide users towards adopting and retaining ergonomic postures.

Traditional biofeedback approaches to inform humans about the risk associated with their posture are traceable to the 1960s. They have mainly employed additional sensory feedback streams conveying information related to the behavior or real-time efforts supported by the user, both for MSD prevention and neurorehabilitation [5–7]. Such biofeedback relies on the monitoring of physiological variables through sensors, the most common of which are electroencephalographs (EEG), for clinical applications [5, 8], electromyographs (EMG) [9], motion capture systems [7, 10–13], force plates [14], and heart rate sensors [15, 16]. Using these signals, one can estimate in real-time variables such as patterns of neural activity [5], joint torques [17–19], and posture [12]. Then, biofeedback consists in generating a sensory stimulation to inform the user with regard to their own behavior, and to trigger warnings in case they exceed a risk or effort threshold. The most common sensory stimulation use visual [20], auditive [20, 21], and somatosensory (usually through vibrators) cues [11, 22–25], or a combination thereof [10, 26]. These traditional approaches been proven to be efficient in modifying human behavior and promoting the adoption of better postures. However, they present two major limitations: (i) they do not allow to guide users towards adopting a precise posture, as this would require a large number of stimulating devices (typically several per joints), and (ii) the need for explicitly interpreting the additional sensory signals of multiple stimulating devices could result in a neural resource allocation problem [27].

In the present paper, we propose to address these limitations by developing an alternative to traditional biofeedback approaches. Specifically, we propose to use an active exoskeleton as an efficient way to (i) measure posture – which requires to identify joints misalignment to be accurate [28, 29] – and (ii) generate sensorimotor stimulation modifying the posture-effort relationship to trigger implicit motor adaptations. This way, the exoskeleton performs both the monitoring and stimulation components of biofeedback, removing the need for additional devices. Importantly, we argue that the sensorimotor stimulation provided by the exoskeleton should not be limited to simple guidance methods, for instance previously used to teach movements [30]. Instead, we propose here to use the exoskeleton to introduce task-relevant variability to promote the adaptation of the central nervous system (CNS) by generating motor errors. This task-relevant characteristic is critical as it is also known that task-irrelevant variability tends to be ignored by humans [31], which could prevent any significant adaptation. This is likely due to the learning of new motor behaviors through the memorization [32] and prediction [33] of movement errors, which is thought to be based on an internal model [34] needing rich and varied input data to robustly control movements [35]. In addition to removing the need for separate and numerous sensory and stimulation devices, such an approach presents the advantage to be highly flexible, as any posture-stimulation mapping can be designed, meaning users could theoretically be guided towards any reachable posture. Furthermore, our approach would only add a stimulation of the somatosensory system through forces that are directly relevant in the task to the normal visual information, which forces are known to be optimizable by the CNS even when unpredictable [36, 37], thereby addressing the risk of a neural resource allocation problem.

However promising, this approach needs to be experimentally tested so as to conclude on its feasibility and usability. Therefore, we designed a reach-to-hold experiment with the following main characteristics (see Figs. 1, 2): (i) the target of the reaching movement was a long vertical rectangle, allowing exploration of various final postures through target redundancy [38], (ii) the movements were in a parasagittal plane, where gravity-related efforts vary significantly, which allowed to differentiate between ergonomic (with lower effort) and non-ergonomic (with higher effort) postures, (iii) participants were exposed to a haptic stimulation, implemented as horizontal vibrations applied by the robot in certain conditions, pushing them outside of the target so as to be task-relevant, and (iv) the vibrations’ amplitude exhibited a minimum (null vibration) that was chosen either above or under the posture each participant nominally preferred. Importantly, when the minimum of the vibrations was above the preferred height, this tended to increase gravity-related efforts (see Fig. 2A), and conversely when the minimum was below. In the rest of the present paper, the combined effects of gravity-related efforts and the height-dependent vibrations applied by the exoskeleton are referred to as the *gravito-vibratory landscape*, the first being ubiquitous and natural while the other is artificially introduced for sensorimotor guidance. The investigations focus on whether participants succeed or fail to adapt by adopting a posture minimizing the intensity of vibrations and/or of gravity-related efforts, and whether they can retain the adapted posture, which would be critical for ergonomics improvement (see Fig. 1C).

**Figure 1:**
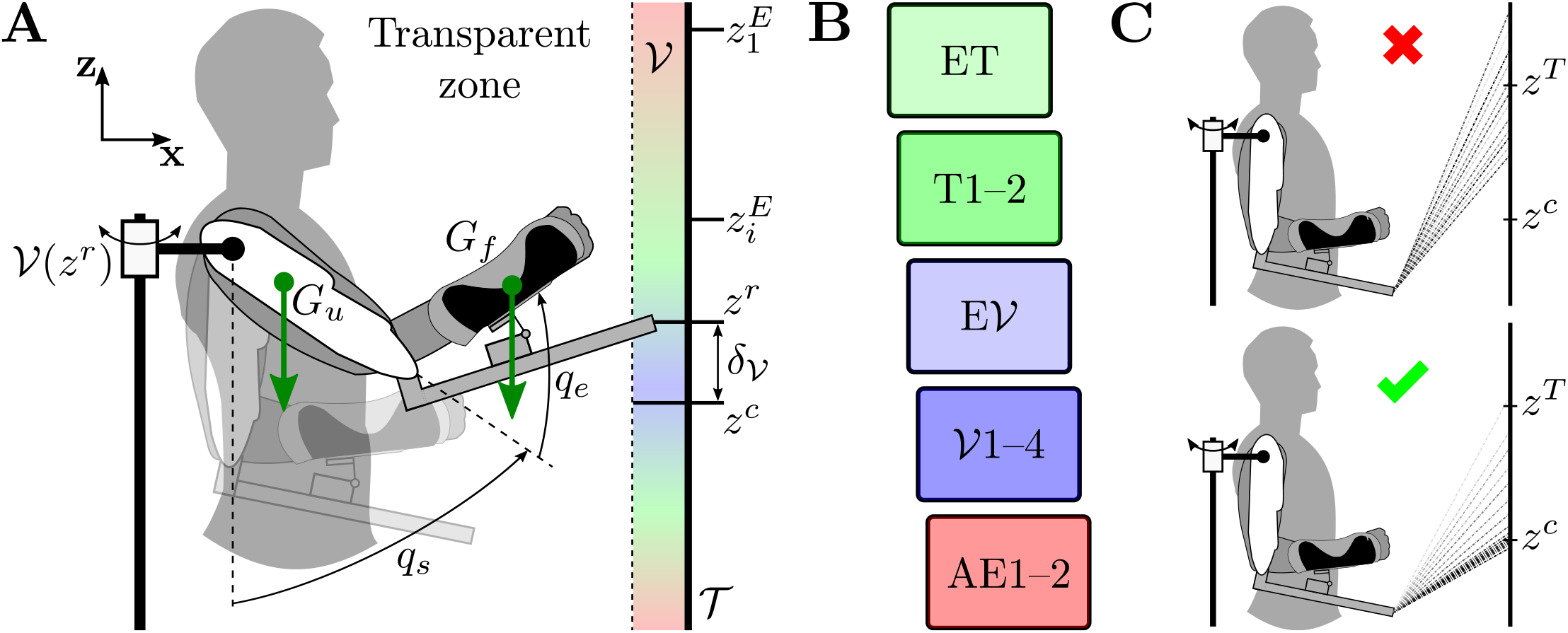
Human-exoskeleton manifold reaching experiment with *gravito-vibratory landscape*. **A**. Participants performed reach-to-hold movements towards a vertical bar 𝒯, which implies a target redundancy as any point on a vertical axis allows to fulfill the task [38]. While maintaining their final posture, participants were subjected to a landscape composed of (i) a gravity component (illustrated by the green arrows), due to the weight of their upper-arm at *G*_*u*_ and forearm at *G*_*f*_, and (ii) a vibratory component𝒱, applied by the exoskeleton depending on the distance between the final height *z*^*r*^ and the center (i.e. cancellation point) of the vibratory landscape *z*^*c*^, referred to as *δ*_*𝒱*_. The center of the landscape is set 20 cm above or below the preferred endpoint height of each participant, resulting in two experimental groups. **B**. The experiment started with an exploration block in transparent mode (ET), where participants were asked to reach towards 10 predefined heights equally spread along the vertical bar 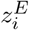, with *i* ∈⟦1, 10⟧, and hold their posture for 3 s. Then, they performed two blocks of reach-to-hold movements in transparent mode (T), i.e. without the vibrations. These were followed by an exploration block similar to ET but with the vibratory landscape (E𝒱 and four blocks with the vibratory landscape activated 𝒱. Finally, two blocks were performed in transparent mode to analyze possible after-effects (AE). **C**. Possible outcomes of the interaction with the landscape from the firsts (light gray) to the lasts (black) trials. Top: the participant fails to converge to *z*^*c*^ and there is no stability in the adopted posture, which remains in the vicinity of *z*^*T*^ (median of the transparent block). Bottom: the participant gradually converges towards *z*^*c*^ and the posture stabilizes around it.

**Figure 2:**
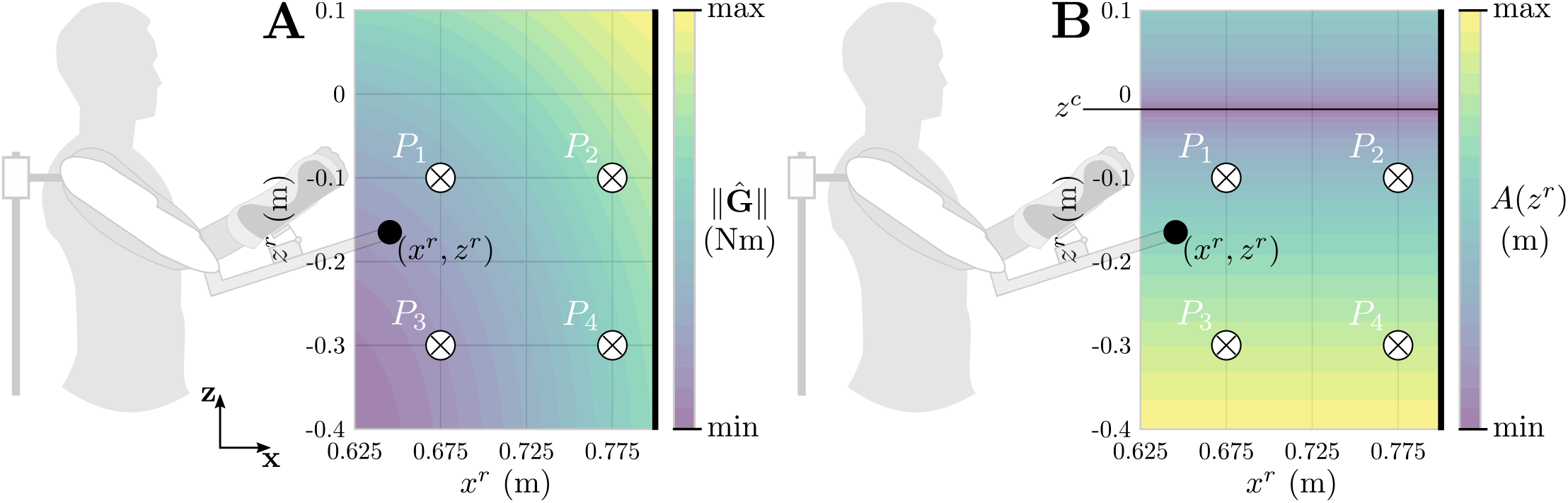
Illustration of the evolution of the two components of the *gravito-vibratory landscape*. Landscapes are computed from 0.4 m below the shoulder to 0.1 m above the shoulder, and from 0.625 m to 0.8 m of robot’s arm extension. *P*_1_–*P*_4_ represent how different end-effector positions can largely affect the effort requested to hold the posture. The black vertical bar on the right represents the target displayed to the participants. **A**. Gravity landscape, with 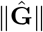 the norm of the gravity torques including the elbow and shoulder. **B**. Vibratory landscape, with *A*(*z*^*r*^) the height-dependent amplitude of the vibration. In this situation, the gradients of both components are of opposite signs, implying that holding a higher posture increases gravity-related efforts but decreases the vibrations magnitude.

## 2 Results

The experiment included three main conditions. First, participants familiarized with the task by performing two blocks of movements with the exoskeleton in transparent mode [39]. This also allowed us to extract the preferred final posture of each participant. Then, they performed four blocks of reach-to-hold trials with a lateral vibration applied by the exoskeleton (𝒱_1_– 𝒱_4_) at the final posture, resulting in a *gravito-vibratory landscape* (see Fig. 1B). Importantly, the amplitude of the vibration was dependent on the endpoint’s height of each participant (see Eq. 1 in Methods), with a null amplitude at the center of the landscape *z*^*c*^ placed at ± 20 cm of the median preferred height of each participant. For half of the participants, whom naturally selected postures low in the target space, *z*^*c*^ was chosen *above* their preferred endpoint height, and it was chosen *below* for the other half of the participants, whom naturally selected postures close to the height of their shoulder. Finally, participants performed two blocks with the transparent exoskeleton so that we could analyze the after-effects (AE) of the exposure to the gravito-vibratory landscape. Importantly, both these components (gravity-related efforts and vibrations) evolved as a function of the final posture to hold (*x*^*r*^, *z*^*r*^) and exhibited a minimum in the reachable space. Furthermore, their gradients were of opposite sign for the *above* group, i.e. reducing the vibrations increased gravity-related efforts, whereas they had the same sign for the *below* group, i.e. reducing vibrations decreased gravity-related efforts.

We first qualitatively analyzed the exploration and optimization of the final posture using representative trajectories throughout the experiment for each group, as summarized in Fig. 3.

**Figure 3:**
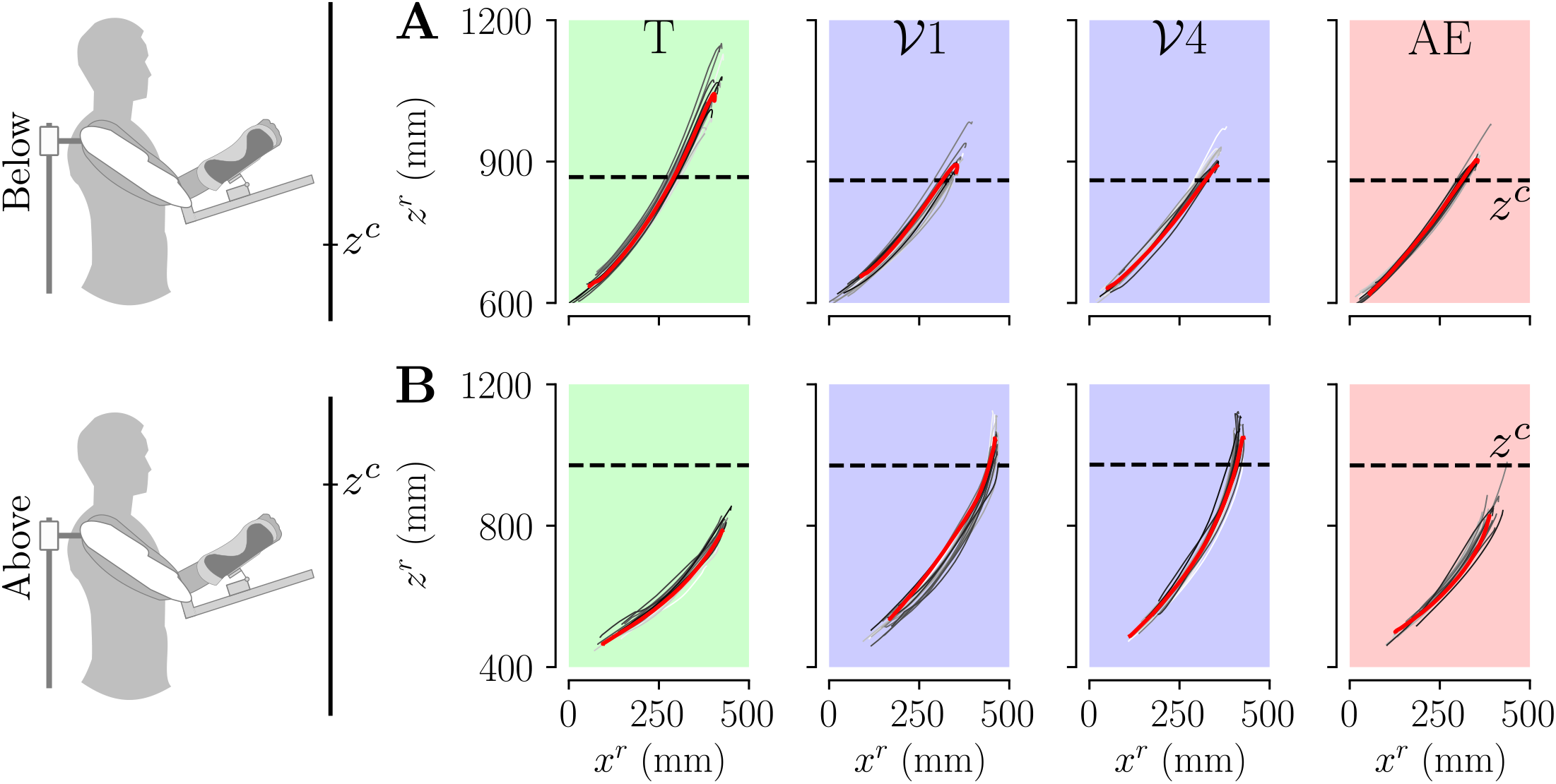
Reaching trajectories of representative participants for each group through the experiment. The red curves represent the trajectories with the median final height (*z*^*r*^) during the block. Horizontal dashed line represents the height at which the amplitude of the vibrations was null. **A**. *Below* group. **B**. *Above* group.

Interestingly, Fig. 3 suggests that both groups adapted their reach-to-hold strategy to minimize the vibrations applied by the exoskeleton at their final posture. Importantly, during the AE blocks, the participant from the *above* group seems to have moved back to their initial strategy while the participant from the *below* group seems to have changed their strategy, retaining their final posture from the blocks with the gravito-vibratory landscape. To better understand how participants explore the landscape, we then extracted “navigation maps” representing the influence of the selected postures on the intensity of each component through the experiment. These analyses are summarized in Fig. 4.

**Figure 4:**
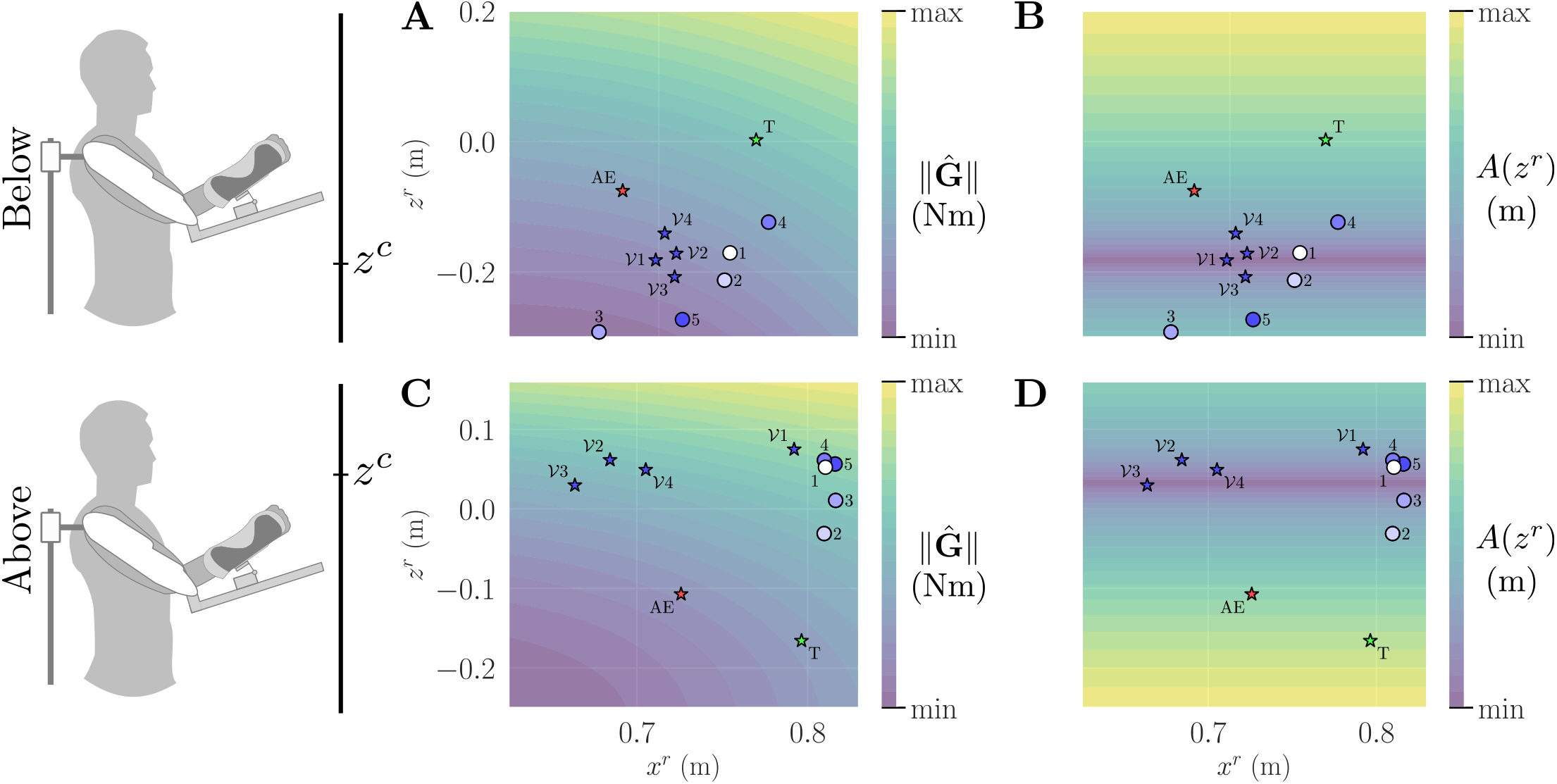
Navigation maps of two representative participants of the *above* and *below* groups. The stars represent the median final position in each block. The disks represent the first five trials in the 𝒱_1_ blocks, showing the initial exploration of the gravito-vibratory landscape. **A**,**C**. Representative navigation of the gravity component for the *below* and *above* group respectively. **B**,**D**. Representative navigation of the vibratory component for the *below* and *above* group respectively.

The navigation maps for the gravity component of the landscape suggest that both groups tended to minimize effort by reducing their arm extension when the landscape was activated. Importantly, for both the *above* and *below* groups, the median final posture in AE seems to retain this reduced arm extension. The navigation of the vibratory component by both representative participants suggests that they tried to minimize the intensity of the vibration. Interestingly, they seem to both succeed to do this through exploration.

Following these qualitative analyses, we quantified the adaptation of the final posture with respect to three main variables: the success rate in each block *ρ*, and the evolutions of *δ*_*𝒱*_ (see Eq. 2) and *δ*_**G**_ (see Eq. 4) throughout the experiment. These two distances, *δ*_*𝒱*_ and *δ*_**G**_, represent how close to the optimum the final posture is with respect to each landscape component. Block-wise analyses of these parameters are summarized in Fig. 5.

**Figure 5:**
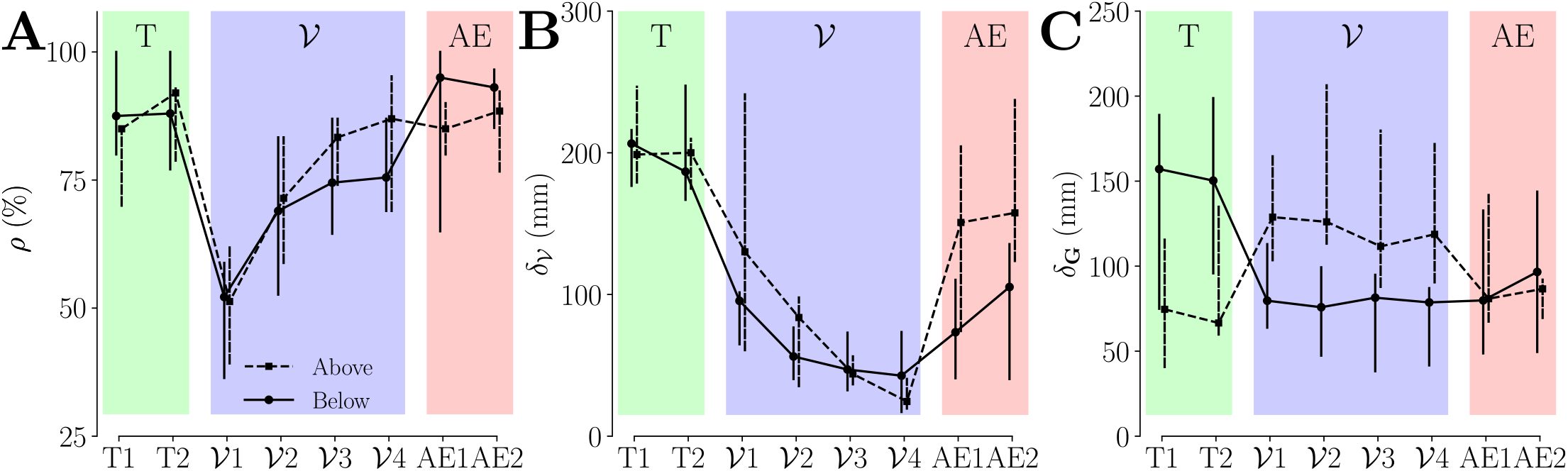
Adaptation of the main interest parameters for each group throughout the experiment. The *above* group is represented with dashed lines and the *below* group with solid lines. Markers represent the median across participants and vertical bars represent the confidence intervals. **A**. Success rate *ρ*. **B**. Distance to the minimum of the vibratory landscape *δ*_*𝒱*_. **C**. Distance to the minimum of the vibratory landscape *δ*_**G**_.

The evolution of the success rate through the blocks 𝒱_1_–𝒱_4_ suggests that participants gradually learned to perform the task, as illustrated by a significant main effect of the block for both the *above* and the *below* group (*χ*^2^ = 3498, *p <* 0.001). Specifically, 𝒱_1_ exhibits a drop in the success rate *ρ* from above 85% to around 50% when compared to the transparent blocks, which was confirmed to be significant for both groups (in both cases: *p <* 0.001, Cohen’s *D >* 3.27). Then, *ρ* gradually increases with learning, as illustrated by the significant differences between 𝒱_1_ and 𝒱_4_ for both groups (in both cases: *p <* 0.001, Cohen’s *D >* 2.48). Interestingly, the performance of the *above* group in 𝒱_4_ was comparable to their performance in the transparent blocks, while the performance of the *below* group was still significantly lower (*p <* 0.001, Cohen’s *D* = 0.984). Finally, the success rate in the AE blocks was similar to the transparent blocks for both groups, which was significantly better than in 𝒱_4_ (in both cases: *p <* 0.001, Cohen’s *D >* 0.47), with a larger and clearer improvement for the *below* group. In sum, the evolution of the success rate throughout the blocks clearly exhibit signs that participants learned how to successfully perform the task. However, this does not allow to conclude regarding their strategy to succeed. In particular, they could either resist to the vibratory landscape through a modulation of their impedance [40], or learn to minimize its intensity through exploring the target space.

To investigate these two possibilities, we analyzed the evolution of *δ*_*𝒱*_, i.e. the distance to the null vibration height, through the blocks 𝒱_1_– 𝒱_4_. For both groups, there was a clear and significant decreasing trend in this distance with practice (*χ*^2^ = 463, *p <* 0.001), which seemed steeper for the *below* group. Specifically, *δ*_*𝒱*_ was significantly smaller in 𝒱_1_ than in the transparent blocks (in both cases: *p <* 0.001, Cohen’s *D >* 1.57). Furthermore, *δ*_V_ tended to decrease between 𝒱_1_ and 𝒱_4_, which decrease was significant for the *above* group (*p <* 0.001, Cohen’s *D* = 0.603). During the AE blocks, participants tended to retain some of the adaptations induced by the vibratory landscape, as illustrated by the significant decrease in *δ*_*𝒱*_ when compared to the transparent blocks (in both cases: *p <* 0.001, Cohen’s *D >* 0.45). However, this retention was not complete as *δ*_𝒱_ significantly increased during AE compared to 𝒱_4_ (in both cases: *p <* 0.001, Cohen’s *D >* 0.72). Remarkably, the *below* group showed a higher retention than the *above* group during AE, as illustrated by significantly lower *δ*_*𝒱*_ (*p <* 0.001, Cohen’s *D* = 0.914). In sum, participants gradually learned to minimize the effects of the vibratory landscape through the four performed blocks. Importantly, the *above* group did not retain much of the learned optimized posture, while the *below* group tended to remain closer to their behavior in 𝒱_4_. Furthermore, the *below* group adapted their behavior to the vibratory landscape faster than the *above* group, although both groups converged to comparable levels of minimization by the end of the four blocks. These important differences between groups suggest that the gravity component of the landscape, and in particular whether the gradients of both landscape components had the same sign, could play a critical role in the adaptability and retention of postures by participants.

Thus, we analyzed the evolution of *δ*_**G**_, i.e. the distance to the minimum of 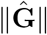 (defined as the norm of the gravity-related efforts, see Eq. 3) within the target space, through the blocks 𝒱_1_– 𝒱_4_. These analyses exhibited a significant main effect of the block on the participants’ final posture (*χ*^2^ = 731, *p <* 0.001). Importantly, due to how they were formed, there was a preexisting difference in *δ*_**G**_ between the *above* group, with lower *δ*_**G**_, and the *below* group, with higher *δ*_**G**_ (*p <* 0.001, Cohen’s *D* = 0.859). Then, as soon as in 𝒱_1_, this difference in distance from the minimum gravity effort was reversed and the *below* group exhibited significantly lower *δ*_**G**_ than the *above* group (*p <* 0.001, Cohen’s *D* = 1.194). Incidentally, the *above* group moved significantly further away from the minimum of efforts, while the *below* group moved closer to it when compared to transparent blocks (in both cases: *p <* 0.001, Cohen’s *D >* 0.95). This effect remained present until the end of the four blocks within the gravito-vibratory landscape, as illustrated by significantly higher *δ*_**G**_ for the *above* than for the *below* group in 𝒱_4_ (*p <* 0.001, Cohen’s *D* = 1.277). Importantly, during the AE blocks, the *above* group moved back to their *δ*_**G**_ of the transparent blocks (no significant difference), which was reminiscent of observations for *δ*_V_. The *below* group partially retained the novel final posture they learned through the interaction with the gravito-vibratory landscape, as illustrated by the small difference between _4_ and AE (*p* = 0.029, Cohen’s *D* = 0.31, small effect) and by a large and significant difference between the transparent blocks and AE (*p <* 0.001, Cohen’s *D* = 0.978). In sum, the similar signs of gradients of the landscape’s components led the *below* group to retain the posture to which they were guided, whereas, as soon as the vibration was removed, the *above* group moved to a less effortful posture.

Importantly, the gravito-vibratory landscape evolves non-linearly in the task space (see Fig. 2). Therefore, although we showed how the posture was retained or not depending on the distance to the minimum effort and minimum vibration, this is not sufficient to conclude regarding the final posture retained by the participants. To quantify more precisely the effects of the gravito-vibratory landscape on the human, we computed the total horizontal distance traveled by the hand after reaching the posture and the average normalized gravity efforts while maintaining it. The normalization of gravity-related efforts was done with respect to the median 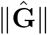 in the transparent blocks, using Eq. 3. The evolution of these two metrics through the experiment is summarized in Fig. 6.

**Figure 6:**
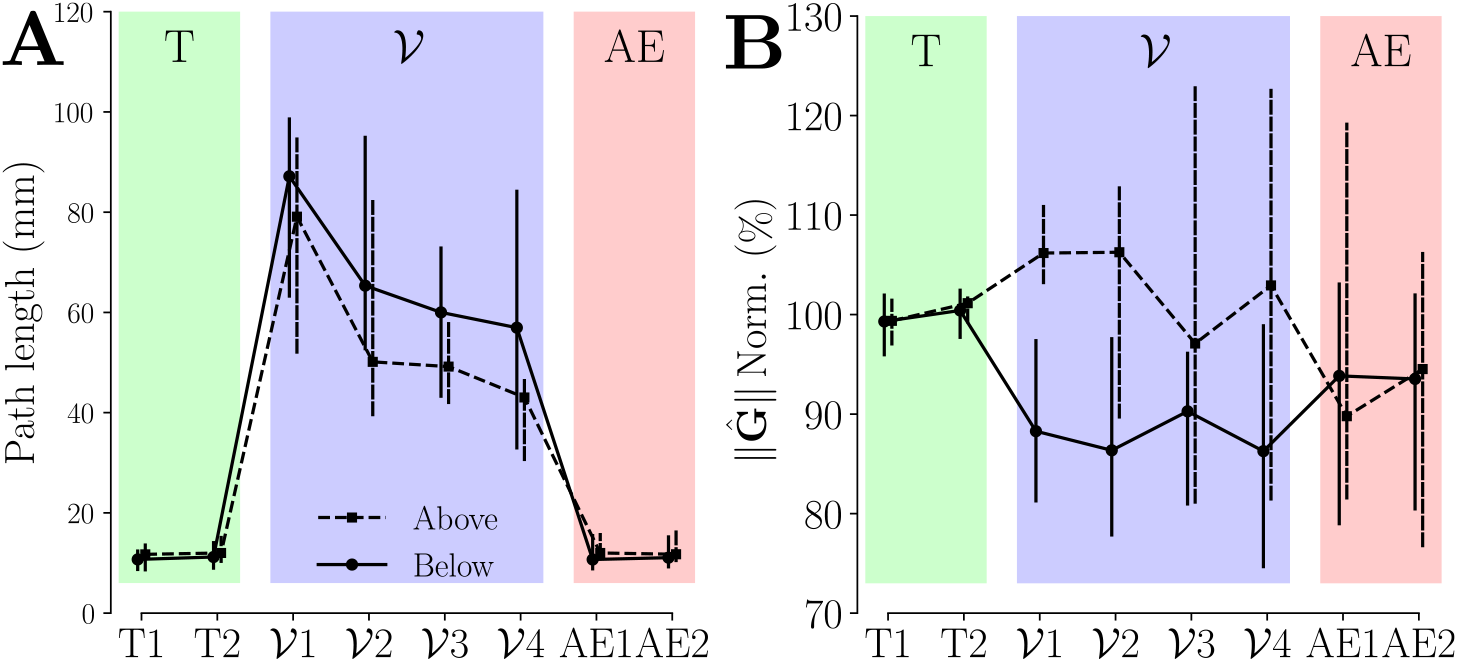
Effects of the gravito-vibratory landscape on the human effort throughout the experiment. The *above* group is represented with dashed lines and the *below* group with solid lines. Markers represent the median across participants and vertical bars represent the confidence intervals. **A**. Total horizontal distance traveled by the hand while maintaining the final posture in the landscape. **B**. Normalized gravity-related efforts averaged while maintaining the final posture in the landscape.

The analysis of the total horizontal displacement of the hand allows to verify how much of the vibration the participants were able to compensate using muscle effort. There was a significant main effect of the block on the participants’ hand horizontal movements (*χ*^2^ = 2039, *p <* 0.001). Specifically, there was only residual horizontal movement in the transparent block, as permitted by passive joints in the human-exoskeleton connection (see Methods). In 𝒱_1_, the horizontal distance traveled by the hand dramatically increased for both groups (in both cases: *p <* 0.001, Cohen’s *D >* 3.36), consistently with observations of *δ*_*𝒱*_ suggesting that they did not find the minimum of the vibration landscape yet. The horizontal movements then decreased with practice to reach a minimum in 𝒱_4_, which was significantly lower than in 𝒱_1_ (in both cases: *p <* 0.001, Cohen’s *D >* 1.29). Interestingly, the *above* group’s hands moved significantly less than the *below* group’s in 𝒱_4_ (*p <* 0.001, Cohen’s *D* = 0.553), suggesting that their posture allowed them to better compensate for the vibrations as no difference was observed in *δ* _𝒱_ in this block. Importantly, for both groups this minimum was still significantly higher than the residual movements observed in both the transparent and AE blocks (in all cases: *p <* 0.001, Cohen’s *D >* 1.75). In sum, participants never completely compensated for the vibrations, which suggests that they only compensated as much vibration as necessary to remain in the target, consistently with a task-dependent minimum intervention hypothesis [41]. This optimal use of the width of the target allows to finalize the analysis of the participant’s behavior with respect to the vibratory component of the landscape.

However, it does not allow to provide a definitive answer regarding the optimization of the gravity component of the landscape. Therefore, we analyzed the evolution of the normalized gravity-related efforts to maintain the final posture throughout the experiment, revealing a significant main effect of the block on the effort provided by the participants (*χ*^2^ = 164, *p <* 0.001). In 𝒱_1_, the effort slightly increased for the *above* group (*p* = 0.015, Cohen’s *D* = 0.353, small effect) and decreased for the *below* group (*p <* 0.001, Cohen’s *D* = 0.494), resulting in divergent effects of the landscape on the effort expended by both groups (*p <* 0.001, Cohen’s *D* = 0.897). There was no clear adaptation of the gravity-related efforts throughout blocks 𝒱_1_ to 𝒱_4_, consistently with results on *δ*_**G**_. Interestingly, the *above* group moved to a posture less effortful in AE when compared to 𝒱_4_ (*p <* 0.001, Cohen’s *D* = 0.476), while no significant difference was observed for the *below* group. Finally, in AE, both groups adopted postures that reduced gravity-related efforts when compared to transparent blocks (in both cases: *p <* 0.022, Cohen’s *D >* 0.26, small effects). In sum, both groups adapted to the gravito-vibratory landscape so as to make task-relevant vibrations smaller than the width of the target, which was observed whether it was beneficial or detrimental to the effort they had to provide. After exposure, despite variability in the results, we could observe that both groups improved their posture in the sense that they reduced the gravity-related efforts.

## 3 Discussion

We investigated whether task-disturbing somatosensory biofeedback applied by a robotic exoskeleton is a feasible method to guide users toward more ergonomic postures, thereby preventing the development of musculoskeletal disorders (MSD). When performing tasks with multiple solutions, humans can sometimes adopt non-ergonomic postures, needlessly putting them at risk of MSD. While this can be due to a number of psychosocial factors, including stress and time constraints [42] or the design of work stations [43, 44], it can also sometimes be the consequence of bad habits and a lack of knowledge about ergonomics [45]. Furthermore, it can also be related to the incapacity of our CNS to consider long-term effects of non-ergonomic postures. In that context, our idea was to design a method leveraging the spontaneous minimization of task-relevant variability in human motor control to teach novel postures to exoskeleton users through posture-dependent mechanical stimulation. Specifically, we designed a *gravito-vibratory landscape* that varied with the user posture and exhibited a minimum for each component within the task target space. The main hypothesis validated by the experiment were twofold. First, the minimization of the landscape intensity through a trial-by-trial gradient descent was possible, allowing participants to find an ergonomic posture, trading off the costs of vibrations and gravity, to perform the task. Second, the participants could retain the novel posture found through exploration if it reduced the effort compared to their preferred posture before exposure, illustrating the benefits of using an exoskeleton to guide the user towards more ergonomic postures.

At the population level, our results clearly show that participants gradually learned to minimize the effects of the vibratory component of the landscape, until the remaining vibrations were sufficiently small to comply with the task constraints. Our vibratory landscape generating large task-relevant variability, it could incentivize the CNS towards performing a “gradient descent” through trials so as to minimize the mechanical effects of the disturbance. The residual vibrations that were not compensated by participants after four blocks of exposure further suggest that the task-relevance of the perturbation is critical for them to adapt, as predicted by the minimum intervention principle [31, 41]. Importantly, our results also showed that participants largely adapted to the gravity component of the landscape within the first block of exposure to vibrations, with small changes in the following blocks, which was clearly faster than the adaptation to the vibratory component. This difference in the time scale of the adaptation is likely due to the omnipresence of gravity in our environment. Specifically, the CNS is thought to plan movements so as to leverage gravity-related efforts [46], as also demonstrated for primates [47] and when interacting with unusual gravity-like torques applied by an exoskeleton [48]. This rapid change in the gravity component of the landscape in reaction to the vibration is particularly important for the prevention of MSD, as gravity-related efforts are known to play a major role in their development. Specifically, ergonomic postures are usually those minimizing fatigue, which often corresponds to minimizing gravity in work conditions, whether it be when carrying loads [49, 50], using screens [51, 52], or when performing highly constrained tasks such as laparascopic surgery [53]. Yet, at the group level, roughly half of our participants naturally selected postures close or above to the shoulder height (*below* group), which implies relatively high, unnecessary, gravity-related effort to maintain it. Interestingly, through the exposure to the gravito-vibratory landscape, we showed that this *below* group rapidly decreased gravity-related efforts in reaction to the task-relevant vibrations, thereby improving their posture. More surprisingly, the *above* group naturally increased gravity-related efforts in a tentative to reduce vibrations, which happened on the same timescale as for the *below* group, showing the higher priority assigned to the successful task completion. These results demonstrate that applying error-generating sensorimotor stimulation to guide a worker wearing an exoskeleton towards novel postures, whether better or worse than their preferred ones, is feasible and that adaptations happen within a relatively small number of trials.

Strikingly, we show contrasted results in the retention of the posture suggested through the gravito-vibratory landscape. Specifically, when the novel final posture obtained through exploration induced higher gravity-related efforts (*above* group), it was clearly not retained by the participants. Conversely, when the minimum of vibrations induced a reduction of these efforts (*below* group), the posture tended to be retained, thereby canceling preexisting differences between groups in the distance to the posture minimizing gravity-related efforts. In both cases, participants showed an improvement of the ergonomics of their preferred posture, as illustrated by the lower norm of gravity-related efforts, suggesting that forcing users to explore a variety of postures, whether better or worse than their preferred one, allows them to minimize the efforts associated with a task. In addition to minimizing the task errors, this finding may be explained in part by the natural movement variability that would allow users to iteratively explore different final postures. Interestingly, there is compelling evidence in movement neuroscience that human motor learning is more efficient when they can explore the task space [54, 55], allowing them to optimize motor behaviors, as made possible by our redundant target. In fact, when asked to perform a novel task, individuals with high levels of task-relevant variability tend to learn faster than those exhibiting lower levels of variability [56]. Therefore, the freedom given to participant in selecting their final posture may contribute to their retention and their improvement of final posture. This illustrates the potential benefits of leveraging this fundamental mechanism of human movement control for ergonomics. However promising, these results would need to be consolidated through longitudinal studies, allowing a longer and repeated exposure as well as long-term retention analyses, which were beyond the scope of the present work. Extending those ideas to whole-body exoskeletons in load-carrying tasks could also be relevant.

When compared to traditional biofeedback methods, our approach solves several important issues. On the sensing side, as it relies on an exoskeleton to measure posture, it does not require any other sensors than those embedded in the robot, which drastically limits the installation and calibration complexity and the required signal processing. This is particularly practical when compared to electromyographic measures of muscle activity [9, 57], or motion capture [7, 10–13] based approaches. However, this means that our method requires knowledge of task-related ergonomic postures, which may not be necessary with other approaches, such as based on recordings of the autonomous nervous system to infer effort [15, 16]. This problem being common to a variety of ergonomic assessments and biofeedback approaches, a large number of tools to address it are available [58], including posture-based scales [59, 60] and more advanced assessments explicitly accounting for effort [61]. On the actuation side, our method only stimulates the sensory streams already involved in performing the task, in particular the somatosensory system at the level of the human-exoskeleton interfaces, which prevents the risk of overloading the brain with sensory information [27]. Furthermore, this stimulation can easily be transposed to any task or posture, without increasing the complexity of the setup, contrary to other biofeedback systems. For instance, when using vibrators as it is commonly done, at least one actuator would be required for each monitored joint to induce a specific posture. The need of an actuated robotic device may be seen as a downside due their cost and the difficulty of their use compared to vibrators, speakers or visual feedback devices (e.g. mixed reality glasses). However, given that exoskeletons are already envisioned as assistive devices for the prevention of MSD [62,63], the proposed approach would therefore require minimal investment when added on top of existing in-field setups.

In case further long-term studies would be conclusive, our method could be applied to an industrial scenario, such as an automotive assembly task with multiple possible postures, using a simple procedure. The first step would be to assess what are the best postures that can be used to perform the task, for instance using existing ergonomic tools [58]. A vibratory landscape with a null intensity near these ideal postures can be implemented, resulting in bad postures being highly penalized. Then, both novice and expert workers can be exposed to this landscape during dummy training sessions, outside of production lines as performance is directly impacted during learning. The exposure sessions can be repeated on a regular basis so as to remind the CNS of the best postures to adopt, thereby preventing the apparition of MSD. Finally, the haptic stimulation can be coupled with an assistance helping to perform the task, for instance by compensating gravity [28, 64] or with adaptive behaviors [65–67], allowing to further reduce the cost of ergonomic postures and paving the path for ergonomic-aware assistance.

## 4 Methods

### 4.1 Participants, materials, and data processing

#### Participants

The experimental protocol was approved by the Université Paris-Saclay’s ethics committee for research (CER-Paris-Saclay-2021-048). A total of *N* = 15 (8 in the *below* and 7 in the *above* group) healthy, right-handed, and naive participants (11 males, and 4 females) participated in the experiment. Their anthropometric characteristics were: age 23.7 ±3.3 years old, height 1.79 ± 0.09 m, and weight 68.80 ± 7.12 kg. Each participant was informed about the experiment, and signed an informed consent form before participating in it.

#### Upper-limb kinematics

We recorded the participant’s upper-limb movements at 100 Hz using an optoelectronic motion capture system (10 Oqus 500+ infrared cameras, Qualisys, Gothenburg, Sweden). Each participant was equipped with 10 mm reflective markers allowing to measure the positions of their upper-arm and forearm, as well as the position of the exoskeleton in real-time (see Fig. 1A for representative positions of the markers). Note that the markers placed on the participants were mainly used to identify the human-exoskeleton joints misalignment in a simple way. In practice, this identification can be performed using other information readily available in the exoskeleton, such as interaction forces [28].

#### ABLE exoskeleton

We used a highly backdriveable robotic upper-limb exoskeleton called *ABLE* [68, 69] to physically interact with the participants (see Fig. 1A). This exoskeleton includes four active degrees of freedom (DoFs), three of which are at the shoulder level (flexion/extension, abduction/adduction, internal/external rotation), and one at the elbow (flexion/extension). This version of ABLE includes ergonomic human-robot physical interfaces to connect the arm and forearm of the participant to the robot [70, 71], which have been shown to reduce unwanted interaction efforts and increase comfort. Furthermore, the exoskeleton is equipped with two force-torque (FT) sensors (1010 digital FT, ATI, sample rate 1 kHz) placed at each human-exoskeleton interface point, which allowed to measure interaction forces to control the robot and improve transparency [39].

The exoskeleton was mostly controlled to be as transparent as possible, which is to follow human movements with a minimal impact on it. The transparent controller relied on the identification and compensation of the exoskeleton’s weight and friction and on the minimization of the interaction efforts using a proportional-integral correction, that was shown to be efficient in previous works [39, 72]. In certain series of trials, when the end-effector of the exoskeleton entered the 𝒱 zone (see Fig. 1B), the motor providing the internal/external rotation of the exoskeleton’s shoulder applied motor perturbations depending on the human posture to generate a vibratory landscape as described below.

#### Gravito-vibratory landscape

The effort landscape, at the end of reach-to-hold movements, generated by the exoskeleton and the dynamics of the human arm was composed of two main components: (i) the vibratory landscape imposed by the exoskeleton, and (ii) the gravity-related efforts to hold the posture.

First, we designed a horizontal vibration with a varying amplitude, which was aimed at perturbing participants when they were holding their final posture. This task-relevant vibration was defined as follows,

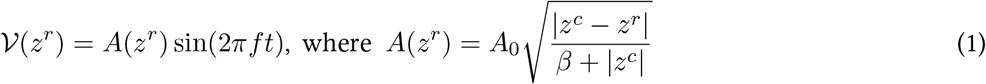

where *A*(*z*^*r*^) is the varying amplitude, *A*_0_ = 9^°^ and *f* = 8 Hz parametrize the amplitude and frequency of the lateral vibration imposed by the exoskeleton, *z*^*r*^ is the vertical coordinate of the exoskeleton’s end-effector in the task space, *z*^*c*^ is the “center” of the landscape, i.e. where the amplitude of the vibration is null, and *β* = 50 was chosen to provide a smooth but sufficiently steep increase of the force field as a function of the distance between *z*^*r*^ and *z*^*c*^. From Eq. 1, one can define the distance to the center of the landscape, which minimizes the effort to provide to laterally stabilize the robot, as follows,

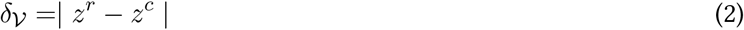

The computation of this distance at the end of each reach-to-hold movement performed by the participant served as one of the main metrics to quantify their adaptation to the vibratory landscape and analyze whether they adopted more ergonomic postures through time, which would imply to observe a gradual decrease of *δ*_*𝒱*_.

Second, we estimated the gravity-related torques 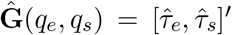 as a function of the participant’s posture using common dynamics,

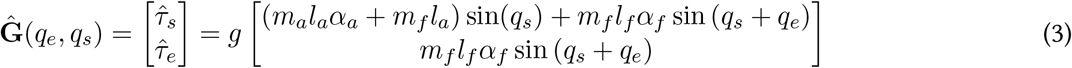

where *q*_*e*_ and *q*_*s*_ are the measured elbow and shoulder joints angles, *m*_*a*_ and *m*_*f*_ are the masses of the arm and forearm, *l*_*a*_ and *l*_*f*_ are the length of the arm and forearm, *α*_*a*_ and *α*_*f*_ are the relative positions of the center of mass of the arm and forearm allowing to obtain *G*_*a*_ and *G*_*f*_ (see Fig. 1B), and 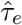 and 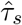 are the estimated elbow and shoulder gravity torques. The length and masses of the segments and the relative positions of center of masses were computed using anthropometric tables [73]. Then, given the vibratory landscape was defined based on the exoskeleton’s end-effector position, we estimated the evolution of gravity-related efforts as a function of the exoskeleton’s posture, thereby defining 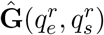 where 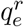 and 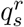 are the exoskeleton’s elbow and shoulder joint angles. Importantly, the human and exoskeleton segments were not aligned due to the passive joints included in the ergonomic interfaces, which implied that 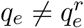 and 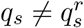. Therefore, to obtain the relationships 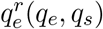 and 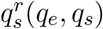, we fitted a 2^*nd*^ order polynomial mapping between the human hand and exoskeleton end-effector position. An illustration of the resulting field, defined using the norm of gravity-related efforts 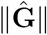 as a function of the exoskeleton end-effector position in the task space (*x*^*r*^, *z*^*r*^), is provided in Fig. 2A. Finally, one can define the distance between the optimal point to reach with respect to the gravity-related efforts and the measured exoskeleton’s end-effector position as follows,

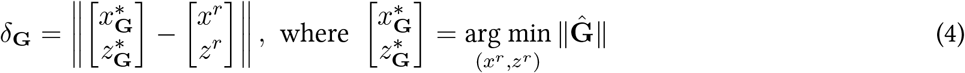

The computation of this distance at the end of each reaching movement performed by the participant will serve as the main metric to quantify their adaptation to the gravity landscape and analyze whether they adopted more ergonomic postures through time, which would imply to observe a gradual decrease of *δ*_**G**_. Values reported in the present paper were obtained by averaging *δ*_*𝒱*_ and *δ*_**G**_ over the last 3 s of each trial, consistently with the validation duration (see Section 4.2).

### 4.2 Manifold reaching task

Participants were asked to perform blocks of reach-to-hold movements in a parasagittal plane using shoulder and elbow flexion/extension movements (see Fig. 1). The start posture of participants was *q*_*s*_ = 0^°^ and *q*_*e*_ = 90^°^, which was controlled in real-time using the motion capture system (see shaded posture in Fig. 1A). The movements were directed towards a 80 cm high and 5 cm large vertical bar displayed on a large screen, thereby defining a manifold in which any point was a valid target. A 2.5 cm blue cursor allowed to show the current height and horizontal position pointed by the exoskeleton’s end-effector to participants as soon as they entered the zone of vibration. Simultaneously, a 3 s countdown was displayed on the screen. This 𝒱 zone’s threshold was set whenever *x*^*r*^ *>* 62.5 cm, with the (**x, z**) frame’s origin placed at the level of the human shoulder. This allowed to ensure that reaching the 𝒱 zone required sufficient movements from the participants while allowing them to easily reach a wide range of vertical positions.

A trial was considered successful if the participant was able to hold its final reaching posture, with the end-effector pointing inside the vertical target, for the whole countdown. In that case, it meant that they were able to compensate for the vibrations. Conversely, a trial was considered failed before the end of the countdown if (i) the participant retracted their arm outside of the vibration zone, (ii) they were not able to maintain the end-effector at a constant height, or (iii) they let the vibrations push them outside of the vertical bar horizontally. The order and requirements of the experimental blocks were as follows (see Fig. 1C),

- **ET – Exploration in transparent mode:** Participants had to perform 10 reach-to-hold movements towards the 10 predefined targets of height 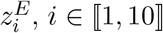 (one movement for each). The trials did not need to be successful.
- **T – Transparent reaching:** Participants had to perform 40 successful reach-to-hold movements towards the vertical bar. The exoskeleton was only controlled in transparent mode, which means that participants were only subject to gravity when maintaining their final posture. This block was performed twice.
- **E𝒱 – Exploration of gravito-vibratory landscape:** Participants had to perform 10 reach-to-hold movements towards the predefined targets as in ET. The trials did not need to be successful.
- **𝒱 – reach-to-hold in gravito-vibratory landscape:** Participants had to perform 20 successful reach-to-hold movements towards the predefined targets. The exoskeleton was controlled in transparent mode outside of the vibration zone. This block was performed four times.
- **AE – After-Effects:** Participants had to perform 40 successful reach-to-hold movements towards the vertical bar. The exoskeleton was only controlled in transparent mode. This block was performed twice.

The exploration blocks allowed the participants to build a prior for each component of the gravito-vibratory landscape, with the objective to facilitate the motor exploration process during the free reaching blocks. To avoid fatigue, breaks of at least 1 minute were taken between each block.

Importantly, *z*^*c*^ was different for each participant, which allowed to normalize the change in final posture height requested to minimize the vibrations. It was set either 20 cm above or 20 cm below the median final height of the participant during the transparent blocks. Two groups of participants were thereby constituted, the “*above*” and the “*below*” groups respectively. The *above* group was constituted of participants that selected final postures below their shoulder height on median, and conversely for the *below* group.

### 4.3 Statistical analyses

We used linear mixed models to assert the main effects of the different conditions and groups on the main studied variables {DV} while accounting for all the trials performed by each participant. These models were as follows,

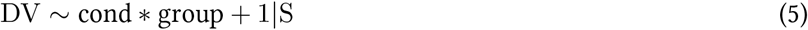

where cond ∈ {T, 𝒱, AE} is the tested condition and group ∈ {downwards, upwards} is the group to which the participant S belongs. This model was then compared to another linear mixed model, without any fixed effect, using a likelihood ratio test, which indicated whether any significant effect of the “cond group” term existed. The linear mixed models analyses were conducted using the *pymer*4 python package [74].

In case of a significant effect, post-hoc comparisons were performed using *t*-tests with a Tukey correction to verify the differences between fixed effects. The level of significance of all the performed tests was set at *p <* 0.05 and we reported Cohen’s *D* as a measure of effect sizes. Whenever the Cohen’s *D* was below 0.4, the effect was flagged as small. The post-hoc analyses were performed using Jasp [75].

## Acknowledgments

This study was funded by the French Agence Nationale de la Recherche (EXOMAN project, grant number ANR-19-CE33-0009).

## Competing interests

The authors declare no competing interests.

## Contribution statement

- Conceptualization – WG, BB, NV, DV
- Methodology – WG, DV
- Software – WG, LQ, DV
- Validation – all authors
- Formal analysis – WG, LQ
- Investigation – WG
- Resources – BB, NV
- Data Curation – WG, LQ
- Visualization – WG, LQ, DV
- Supervision – BB, NV, DV
- Project administration – BB, NV
- Funding acquisition – BB, NV
- Writing - Original Draft – WG, DV
- Writing - Review & Editing – all authors

## Notes

### Competing Interest Statement

The authors have declared no competing interest.

## References

[1] J. Mehrholz, A. Pollock, M. Pohl, J. Kugler, and B. Elsner, “Systematic review with network meta-analysis of randomized controlled trials of robotic-assisted arm training for improving activities of daily living and upper limb function after stroke,” Journal of NeuroEngineering and Rehabilitation, vol. 17, pp. 1–14, jun 2020.

[2] M. A. Nussbaum, B. D. Lowe, M. de Looze, C. Harris-Adamson, and M. Smets, “An introduction to the special issue on occupational exoskeletons,” IISE Transactions on Occupational Ergonomics and Human Factors, vol. 7, pp. 153–162, oct 2019.

[3] T. Luger, M. Bär, R. Seibt, M. A. Rieger, and B. Steinhilber, “Using a back exoskeleton during industrial and functional tasks– effects on muscle activity, posture, performance, usability, and wearer discomfort in a laboratory trial,” Human Factors: The Journal of the Human Factors and Ergonomics Society, vol. 65, pp. 5–21, Apr. 2021.

[4] S. E. Kranenborg, C. Greve, M. F. Reneman, and C. C. Roossien, “Side-effects and adverse events of a shoulder- and back-support exoskeleton in workers: a systematic review,” Applied Ergonomics, vol. 111, p. 104042, Sept. 2023.

[5] S. Horowitz, “Biofeedback applications: a survey of clinical research,” Alternative & complementary therapies, vol. 12, no. 6, pp. 275–281, 2006.

[6] O. M. Giggins, U. Persson, and B. Caulfield, “Biofeedback in rehabilitation,” Journal of NeuroEngineering and Rehabilitation, vol. 10, no. 1, p. 60, 2013.

[7] D. R. Martins, S. M. Cerqueira, and C. P. Santos, “Combining inertial-based ergonomic assessment with biofeedback for posture correction: A narrative review,” Computers & Industrial Engineering, vol. 190, p. 110037, Apr. 2024.

[8] T. M. Sokhadze, R. L. Cannon, and D. L. Trudeau, “EEG biofeedback as a treatment for substance use disorders: review, rating of efficacy, and recommendations for further research,” Applied Psychophysiology and Biofeedback, vol. 33, pp. 1–28, Jan. 2008.

[9] P. Madeleine, P. Vedsted, A. K. Blangsted, G. Sjoegaard, and K. Soegaard, “Effects of electromyographic and mechanomyographic biofeedback on upper trapezius muscle activity during standardized computer work,” Ergonomics, vol. 49, pp. 921–933, Aug. 2006.

[10] N. Vignais, M. Miezal, G. Bleser, K. Mura, D. Gorecky, and F. Marin, “Innovative system for real-time ergonomic feedback in industrial manufacturing,” Applied Ergonomics, vol. 44, pp. 566–574, jul 2013.

[11] S. M. Cerqueira, A. F. D. Silva, and C. P. Santos, “Smart vest for real-time postural biofeedback and ergonomic risk assessment,” IEEE Access, vol. 8, pp. 107583–107592, 2020.

[12] C. M. Lind, “A rapid review on the effectiveness and use of wearable biofeedback motion capture systems in ergonomics to mitigate adverse postures and movements of the upper body,” Sensors, vol. 24, p. 3345, May 2024.

[13] A. Lindegård, A. Grimby-Ekman, J. Wahlström, and E. Gustafsson, “Can biofeedback training in combination with ergonomic information reduce pain among young adult computer users with neck and upper extremity symptoms? A randomized controlled intervention study,” Applied Ergonomics, vol. 114, p. 104155, Jan. 2024.

[14] W. Kim, M. Lorenzini, K. Kapicioglu, and A. Ajoudani, “ErgoTac: a tactile feedback interface for improving human ergonomics in workplaces,” IEEE Robotics and Automation Letters, vol. 3, pp. 4179–4186, oct 2018.

[15] T.-W. Shen, T. Hsiao, Y.-T. Liu, and T.-Y. He, “An ear-lead ECG based smart sensor system with voice biofeedback for daily activity monitoring,” in TENCON - IEEE Region 10 Conference, pp. 1–6, Nov. 2008.

[16] A. P. Sutarto, M. N. A. Wahab, and N. M. Zin, “Heart rate variability (HRV) biofeedback: a new training approach for operator’s performance enhancement,” Journal of industrial engineering and management, vol. 3, no. 1, pp. 176–198, 2010.

[17] K. Kiguchi and Y. Hayashi, “An EMG-based control for an upper-limb power-assist exoskeleton robot,” IEEE Transactions on Systems, Man, and Cybernetics, Part B (Cybernetics), vol. 42, pp. 1064–1071, Aug. 2012.

[18] T. Teramae, T. Noda, and J. Morimoto, “EMG-based model predictive control for physical human–robot interaction: application for assist-as-needed control,” IEEE Robotics and Automation Letters, vol. 3, pp. 210–217, jan 2018.

[19] L. Quesada, D. Verdel, O. Bruneau, B. Berret, M.-A. Amorim, and N. Vignais, “EMG-to-torque models for exoskeleton assistance: a framework for the evaluation of in situ calibration,” bioRχiv, Jan. 2024.

[20] N. Hasegawa, K. Takeda, M. Sakuma, H. Mani, H. Maejima, and T. Asaka, “Learning effects of dynamic postural control by auditory biofeedback versus visual biofeedback training,” Gait & Posture, vol. 58, pp. 188–193, Oct. 2017.

[21] W.-G. Yoo and S.-Y. Park, “Effects of posture-related auditory cueing (PAC) program on muscles activities and kinematics of the neck and trunk during computer work,” Work, vol. 50, no. 2, pp. 187–191, 2015.

[22] C. Ho, H. Z. Tan, and C. Spence, “The differential effect of vibrotactile and auditory cues on visual spatial attention,” Ergonomics, vol. 49, pp. 724–738, jun 2006.

[23] S. Schatzle, T. Ende, T. Wusthoff, and C. Preusche, “VibroTac: An ergonomic and versatile usable vibrotactile feedback device,” in International Symposium in Robot and Human Interactive Communication, sep 2010.

[24] M. Lorenzini, J. M. Gandarias, L. Fortini, W. Kim, and A. Ajoudani, “Ergotac-belt: anticipatory vibrotactile feedback to lead centre of pressure during walking,” in IEEE RAS/EMBS International Conference for Biomedical Robotics and Biomechatronics (BioRob), pp. 01–06, Aug. 2022.

[25] N. A. Kelly, A. Althubaiti, A. D. Katapadi, A. G. Smith, S. C. Nyirjesy, J. H. Yu, A. J. Onwuka, and T. Chiang, “Association of vibrotactile biofeedback with reduced ergonomic risk for surgeons during tonsillectomy,” JAMA Otolaryngology–Head & Neck Surgery, vol. 149, p. 397, May 2023.

[26] M. M. Glumm, K. L. Kehring, and T. L. White, “Effects of visual and auditory cues about threat location on target acquisition and attention to auditory communications,” Proceedings of the Human Factors and Ergonomics Society Annual Meeting, vol. 49, pp. 347–351, sep 2005.

[27] G. Dominijanni, S. Shokur, G. Salvietti, S. Buehler, E. Palmerini, S. Rossi, F. De Vignemont, A. d’Avella, T. R. Makin, D. Prattichizzo, and S. Micera, “The neural resource allocation problem when enhancing human bodies with extra robotic limbs,” Nature Machine Intelligence, vol. 3, pp. 850–860, Oct. 2021.

[28] D. Verdel, S. Bastide, N. Vignais, O. Bruneau, and B. Berret, “Human weight compensation with a backdrivable upper-limb exoskeleton: identification and control,” Frontiers in Bioengineering and Biotechnology, vol. 9, pp. 1–16, jan 2022.

[29] R. Mallat, M. Khalil, G. Venture, V. Bonnet, and S. Mohammed, “Human-exoskeleton joint misalignment: a systematic review,” in 2019 Fifth International Conference on Advances in Biomedical Engineering (ICABME), IEEE, oct 2019.

[30] D. J. Reinkensmeyer and J. L. Patton, “Can robots help the learning of skilled actions?,” Exercise and Sport Sciences Reviews, vol. 37, pp. 43–51, Jan. 2009.

[31] E. Todorov and M. Jordan, “A minimal intervention principle for coordinated movement,” Advances in neural information processing systems, vol. 15, 2002.

[32] D. J. Herzfeld, P. A. Vaswani, M. K. Marko, and R. Shadmehr, “A memory of errors in sensorimotor learning,” Science, vol. 345, pp. 1349–1353, Sept. 2014.

[33] K. Takiyama, M. Hirashima, and D. Nozaki, “Prospective errors determine motor learning,” Nature Communications, vol. 6, Jan. 2015.

[34] M. Kawato, “Internal models for motor control and trajectory planning,” Current Opinion in Neurobiology, vol. 9, pp. 718–727, ec 1999.

[35] M. Bar, Predictions in the brain: Using our past to generate a future. Oxford University Press, 2011.

[36] E. Burdet, R. Osu, D. W. Franklin, T. Yoshioka, T. E. Milner, and M. Kawato, “A method for measuring endpoint stiffness during multi-joint arm movements,” Journal of Biomechanics, vol. 33, pp. 1705–1709, Dec. 2000.

[37] B. Berret, D. Verdel, E. Burdet, and F. Jean, “Co-contraction embodies uncertainty: an optimal feedforward strategy for robust motor control,” PLOS Computational Biology, vol. 20, no. 11, p. e1012598, 2024.

[38] B. Berret, E. Chiovetto, F. Nori, and T. Pozzo, “Manifold reaching paradigm: how do we handle target redundancy?,” Journal of Neurophysiology, vol. 106, pp. 2086–2102, oct 2011.

[39] D. Verdel, A. Farr, T. Devienne, N. Vignais, B. Berret, and O. Bruneau, “Human movement modifications induced by different levels of transparency of an active upper limb exoskeleton,” Frontiers in Robotics and AI, vol. 11, Jan. 2024.

[40] E. Burdet, R. Osu, D. W. Franklin, T. E. Milner, and M. Kawato, “The central nervous system stabilizes unstable dynamics by learning optimal impedance,” Nature, vol. 414, pp. 446–449, Nov. 2001.

[41] A. Takagi, D. Verdel, and E. Burdet, “Intermittent movement control emerges from information-based planning,” bioRχiv, Mar. 2025.

[42] C. Deeney and L. O’Sullivan, “Work related psychosocial risks and musculoskeletal disorders: potential risk factors, causation and evaluation methods,” Work, vol. 34, no. 2, pp. 239–248, 2009.

[43] E. R. Vieira and S. Kumar, “Working postures: a literature review,” Journal of Occupational Rehabilitation, vol. 14, pp. 143–159, June 2004.

[44] N. Jaffar, A. H. Abdul-Tharim, I. F. Mohd-Kamar, and N. S. Lop, “A literature review of ergonomics risk factors in construction industry,” Procedia Engineering, vol. 20, pp. 89–97, 2011.

[45] Y. Salik and A. Özcan, “Work-related musculoskeletal disorders: a survey of physical therapists in Izmir-Turkey,” BMC Musculoskeletal Disorders, vol. 5, Aug. 2004.

[46] O. White, J. Gaveau, L. Bringoux, and F. Crevecoeur, “The gravitational imprint on sensorimotor planning and control,” Journal of Neurophysiology, vol. 124, pp. 4–19, jul 2020.

[47] J. Gaveau, S. Grospretre, B. Berret, D. E. Angelaki, and C. Papaxanthis, “A cross-species neural integration of gravity for motor optimization,” Science Advances, vol. 7, p. eabf7800, apr 2021.

[48] D. Verdel, S. Bastide, F. Geffard, O. Bruneau, N. Vignais, and B. Berret, “Reoptimization of single-joint motor patterns to non-earth gravity torques induced by a robotic exoskeleton,” iScience, vol. 26, p. 108350, Nov. 2023.

[49] D. Abe, S. Muraki, and A. Yasukouchi, “Ergonomic effects of load carriage on the upper and lower back on metabolic energy cost of walking,” Applied Ergonomics, vol. 39, pp. 392–398, May 2008.

[50] A. Koltan, “An ergonomics approach model to prevention of occupational musculoskeletal injuries,” International Journal of Occupational Safety and Ergonomics, vol. 15, pp. 113–124, Jan. 2009.

[51] L. Straker, R. Skoss, A. Burnett, and R. Burgess-Limerick, “Effect of visual display height on modelled upper and lower cervical gravitational moment, muscle capacity and relative strain,” Ergonomics, vol. 52, pp. 204–221, Feb. 2009.

[52] W. Tapanya, R. Puntumetakul, M. Swangnetr Neubert, and R. Boucaut, “Influence of neck flexion angle on gravitational moment and neck muscle activity when using a smartphone while standing,” Ergonomics, vol. 64, pp. 900–911, Jan. 2021.

[53] S. Liu, D. Hemming, R. B. Luo, J. Reynolds, J. C. Delong, B. J. Sandler, G. R. Jacobsen, and S. Horgan, “Solving the surgeon ergonomic crisis with surgical exosuit,” Surgical Endoscopy, vol. 32, pp. 236–244, June 2017.

[54] A. K. Dhawale, M. A. Smith, and B.P. Ölveczky, “The role of variability in motor learning,” Annual Review of Neuroscience, vol. 40, pp. 479–498, July 2017.

[55] O. Ozen, K. A. Buetler, and L. Marchal-Crespo, “Promoting motor variability during robotic assistance enhances motor learning of dynamic tasks,” Frontiers in Neuroscience, vol. 14, Feb. 2021.

[56] H. G. Wu, Y. R. Miyamoto, L. N. G. Castro, B.P. Ölveczky, and M. A. Smith, “Temporal structure of motor variability is dynamically regulated and predicts motor learning ability,” Nature Neuroscience, vol. 17, pp. 312–321, jan 2014.

[57] L. Quesada, D. Verdel, O. Bruneau, B. Berret, M.-A. Amorim, and N. Vignais, “EMG feature extraction and muscle selection for continuous upper limb movement regression,” Biomedical Signal Processing and Control, vol. 103, p. 107323, May 2025.

[58] P. G. Dempsey, R. W. McGorry, and W. S. Maynard, “A survey of tools and methods used by certified professional ergonomists,” Applied Ergonomics, vol. 36, pp. 489–503, July 2005.

[59] L. McAtamney and E. N. Corlett, “RULA: a survey method for the investigation of work-related upper limb disorders,” Applied Ergonomics, vol. 24, pp. 91–99, apr 1993.

[60] S. Hignett and L. McAtamney, “Rapid entire body assessment (REBA),” Applied Ergonomics, vol. 31, pp. 201–205, apr 2000.

[61] P. Maurice, V. Padois, Y. Measson, and P. Bidaud, “Experimental assessment of the quality of ergonomic indicators for dynamic systems computed using a digital human model,” International Journal of Human Factors Modelling and Simulation, vol. 5, no. 3, p. 190, 2016.

[62] M. P. de Looze, T. Bosch, F. Krause, K. S. Stadler, and L. W. O’Sullivan, “Exoskeletons for industrial application and their potential effects on physical work load,” Ergonomics, vol. 59, pp. 671–681, May 2016.

[63] J. Theurel and K. Desbrosses, “Occupational exoskeletons: overview of their benefits and limitations in preventing work-related musculoskeletal disorders,” IISE Transactions on Occupational Ergonomics and Human Factors, vol. 7, pp. 264–280, July 2019.

[64] F. Just, Özhan Özen, S. Tortora, V. Klamroth-Marganska, R. Riener, and G. Rauter, “Human arm weight compensation in rehabilitation robotics: efficacy of three distinct methods,” Journal of NeuroEngineering and Rehabilitation, vol. 17, no. 1, 2020.

[65] T. Proietti, V. Crocher, A. Roby-Brami, and N. Jarrasse, “Upper-Limb Robotic Exoskeletons for Neurorehabilitation: A Review on Control Strategies,” IEEE Reviews in Biomedical Engineering, vol. 9, pp. 4–14, 2016.

[66] S. Dalla Gasperina, L. Roveda, A. Pedrocchi, F. Braghin, and M. Gandolla, “Review on patient-cooperative control strategies for upper-limb rehabilitation exoskeletons,” Frontiers in Robotics and AI, vol. 8, Dec. 2021.

[67] Y. Li, A. Sena, Z. Wang, X. Xing, J. Babič, E. van Asseldonk, and E. Burdet, “A review on interaction control for contact robots through intent detection,” Progress in Biomedical Engineering, vol. 4, p. 032004, jul 2022.

[68] P. Garrec, J.-P. Friconneau, Y. Méasson, and Y. Perrot, “ABLE, an Innovative Transparent Exoskeleton for the Upper-Limb,” IEEE/RSJ International Conference on Intelligent Robots and Systems (IROS), pp. 1483–1488, Sept. 2008.

[69] P. Garrec, “Screw and Cable Acutators (SCS) and Their Applications to Force Feedback Teleoperation, Exoskeleton and Anthropomorphic Robotics,” Robotics 2010 Current and Future Challenges, pp. 167–191, 2010.

[70] D. Verdel, G. Sahm, S. Bastide, O. Bruneau, B. Berret, and N. Vignais, “Influence of the physical interface on the quality of human–exoskeleton interaction,” IEEE Transactions on Human-Machine Systems, vol. 53, no. 1, pp. 44–53, 2022.

[71] D. Verdel, G. Sahm, O. Bruneau, B. Berret, and N. Vignais, “A trade-off between complexity and interaction quality for upper limb exoskeleton interfaces,” Sensors, vol. 23, p. 4122, apr 2023.

[72] D. Verdel, S. Bastide, N. Vignais, O. Bruneau, and B. Berret, “An identification-based method improving the transparency of a robotic upper-limb exoskeleton,” Robotica, vol. 39, pp. 1711–1728, Sept. 2021.

[73] D. A. Winter, Biomechanics and motor control of human movement. New York: John Wiley and Sons, second ed., 1990.

[74] E. Jolly, “Pymer4: connecting R and Python for linear mixed modeling,” Journal of Open Source Software, vol. 3, p. 862, Nov. 2018.

[75] JASP Team, “JASP (Version 0.19.3)[Computer software],” 2025.

